# The Effect of the Glossary on the Reliability and Performance of the Reading the Mind in the Eyes Test (RMET)

**DOI:** 10.1101/2020.03.25.007575

**Authors:** José Toloza-Muñoz, Jean González-Mendoza, Ramón D. Castillo, Diego Morales-Bader

**Author notes:** These authors contributed equally to this work. Universidad de Talca, Avenida Lircay s/n, Talca Chile, CP 3460000.

## Abstract

The Reading the Mind in the Eyes Test (RMET) is used to measure high-level Theory of Mind. RMET consists of images of the regions surrounding the eyes and a glossary of terms that defines words associated with the gazes depicted in the images. People must identify the meaning associated with each gaze and can consult the glossary as they respond. The results indicate that typically developed adults perform better than adults with neurodevelopmental disorders. However, the evidence regarding the validity and reliability of the test is contradictory. This study evaluated the effect of the glossary on the performance, internal consistency, and temporal stability of the test. A total of 89 undergraduate students were randomly assigned to three conditions. The first group used the original glossary (Ori-G). The second group developed a self-generated glossary of gazes (Self-G). Finally, the third group developed a glossary that did not define gazes, but unrelated words instead (No-G). The test was administered before and after participants drew a randomly assigned image as a secondary task. The findings show that the number of correct answers was similar among the three conditions before and after the secondary task. However, the Self-G and No-G groups took less time to finish the test. The type of glossary affected the consistency and stability of the test. In our case, the Self-G condition made the responses faster, more consistent, and more stable. The results are discussed in terms of levels of processing and the detection of mental states based on gazes.

## Introduction

Theory of Mind (ToM) is the ability of human beings, as well as some primates, to attribute, understand and interpret our own or other people’s mental states, thoughts, beliefs, desires, and intentions [1, 2]. The study of ToM has been important in understanding individual differences in social adaptability, which is diminished in people with certain developmental and psychiatric disorders [3, 4].

The ToM is fully developed in people with typical development after around 5 years of age [5]. That means that most adults with normal development should exhibit a well-developed ToM, especially when they do not have deficits in cognitive functions.

A way of measuring ToM in people with high-level cognitive functions is by assessing their ability to capture the mental and emotional state of others by observing their gaze. Baron-Cohen and collaborators created the Reading the Mind in the Eyes Test (RMET) due to the need for an appropriate test to detect variability in the ToM ability among adults with typical development [2, 6]. The instrument consists of 36 photographs showing the gaze of men and women expressing a feeling or thought. Each photograph has four possible answers that appear on the screen. People must choose the most appropriate one.

RMET is considered a more advanced test as it values the complex emotional aspects that arise in social interactions, in addition to the fact that the subject must evaluate another person’s point of view based on an aspect of their face such as the regions that surround the eyes [2]. However, since intentions must be interpreted from what facial expressions reflect, this test could be considered to measure emotion recognition rather than mental state or intentionality [7].

The RMET has been able to capture differences in ToM capacity between men and women, where women score higher [2, 6, 8–10]. This test has also been able to find differences between clinical populations and control groups [9, 11–13]. However, the RMET does not work well when discriminating between individuals with average Theory of Mind skills and those with high skills, as most items have poor discrimination capacity [14].

In some studies, the RMET seems to be reliable in its temporal stability [10, 15–17]. However, some studies have carried out reliability reviews and obtained varied results regarding their internal consistency. The average internal consistency of the studies reviewed is 0.64 *±* 0.12. Taking into account all the calculation methods used (Cronbach’s Alpha, Split-Half, KR20, Ordinal Cronbach’s Alpha, Ordinal Omega, Maximal Weighted Internal Consistency Reliability for the Unidimensional Model), the values fluctuate between a minimum of .37 and a maximum of .77 [8, 10, 14, 17–29]. Müller and Gmünder [24] point out that tests with dichotomous scores usually have lower alpha coefficients than those using Likert scales. However, this does not explain why there is such variability in the internal consistency reported by the prior studies.

The RMET has also reported differences in the average scores obtained by the aforementioned studies. The variability in scores could be explained by verbal IQ level, which contributes significantly to the performance variation in the test [30, 31]. Considering the importance of the subjects’ level of verbal processing on test variability, we believe that the use of the glossary of terms deserves to be studied. Experimental manipulations in which the participants carried out in-depth processing has enabled better performance in terms of accuracy and speed, that is, better retrieval of learned information with less material forgotten [32, 33]. We wonder whether the experimental manipulation of this glossary leads to differences in people’s performance, internal consistency and the temporal stability of the RMET. Therefore, we propose generating two processing conditions that affect the participant’s performance and test reliability (stability and internal consistency). To that end, we introduced variations in the glossary as follows: In one condition, the participant had to generate a list of synonyms or definitions for the words of the original glossary with their own terms that they collected on the Internet, as a Self-Generated Glossary (Self-G). In the second condition, the participant did not have a glossary of words related to the gazes (No-G), although they generated a glossary of neutral words that referred to different meanings. Finally in the third condition participants had to read and learn the definitions of the original glossary (Ori-G). We expected people’s performance to be better and more consistent when they had to craft their own glossary of terms than when they read the glossary of the original test (Ori-G). Finally, we hypothesized that Self-G and Ori-G conditions would have better performance, greater internal consistency, and more temporal stability than the condition in which the participant had no glossary available (No-G).

## Materials and methods

### Participants

A total of 89 undergraduate students (21 men and 68 women) participated, with ages ranging from 18 to 29 years old (M = 21.34, SD = 2.01). The participants were divided into the following groups: Ori-G (N = 30), Self-G (N = 29), and No-G (N = 30). All participants were presented with informed consent before conducting the experiment validated by the University of Talca’s Ethics Committee (FONDECYT 1161533).

### Materials and Procedure

We used Baron-Cohen et al. “Reading the Mind in the Eyes Test” [2] adapted to the Spanish language by Serrano and Allegri [34]. This test contains 37 images (one for practice and thirty-six for evaluation), which only show the regions that surround the eyes. For each of the images, participants see four possible answer options, from which they must choose only one. Participants also have a glossary containing the words that appear next to the images, so that participants can consult it as many times as they want if they have doubts regarding the meaning of the words.

The experiment was divided into four phases. The first phase varied according to each condition and was 25 minutes long. In the Ori-G condition, participants reviewed the list of words and definitions included in the RMET. In the Self-G condition, participants read the original list of words; but they had to create at least one synonym for each word and one sentence for each synonym. Both groups were allowed to revisit the glossary during the experiment. In the No-G condition, participants read a list of words unrelated to the terms in the test and they had to create at least one synonym for each word and one sentence for each synonym. They were not allowed to revisit the glossary during the experiment.

In phase two, participants answered the Reading the Mind in the Eyes Test that had been set up on a computer with E-prime 3.0 software. This program displayed the instructions and the sample image. The program then randomly showed the 36 images that constitute the RMET, recording the response and the total time the person used to respond to each image. The instructions were as follows:

*“In the next task, on the computer screen, a series of images will appear which correspond to the regions that surround the eyes of different people. For each image, select the word that best describes what the person in the picture thinks or feels by pressing the corresponding number on the numeric keypad. It may seem to you that more than one word is applicable to an image, but please choose only one word, the word that you consider as the most appropriate. Before making your choice, make sure you have read all 4 words. You should try to perform this task as quickly as possible*.*”* For the Ori-G and Self-G conditions, the following statement appeared: *“If you really don’t know the meaning of a word, you can look it up in the glossary*.*”* Thus, participants in the Ori-G and Self-G conditions could use the glossary of words during the test. For the No-G condition, the following statement appears: *“If you really don’t know the meaning of a word, you should try to guess which one might be right*.*”*

In phase three, participants performed a secondary task, which consisted of drawing a randomly assigned image (car, train, boat, plane, or bicycle). Participants had three minutes to complete the drawing. Finally, in phase four, the Reading the Mind in the Eyes Test was applied again, using the same method as the first application.

### Analysis

We calculated the internal consistency for the entire test and all three conditions using Cronbach’s Alpha and the Kuder–Richardson 20 (KR-20) formula, which is a special case of Cronbach’s Alpha used for calculating dichotomous items. To assess the temporal stability of the test, we calculated the Spearman-Brown coefficient. Additionally, we evaluated the consistency between the test and retest with the interclass correlation coefficient using the average fixed raters (ICC3k). The ICC is a measure that evaluates the reproducibility of repeated measurements in the same population. Scores equal to or greater than .60 are considered acceptable for clinical use [35]. In addition to the ICC, the distribution of score differences were analyzed with Bland-Altman plots.

To assess whether there were differences between the three conditions and between the test and the retest, we performed a mixed ANOVA with a 3×2 design with the averages of correct answers and the averages of the total test time.

## Results

### Reliability

The internal consistency of the test was low for the three conditions (Cronbach’s *α <* .70) and for the total test (Cronbach’s *α* test = .29, Cronbach’s *α* retest = .51), both in the test and in the retest. The test under the Self-G condition obtained greater internal consistency, while the test with the Ori-G obtained the lowest consistency in both the test and the retest. KR-20 was also calculated, but no changes in trends were observed when compared to Cronbach’s Alpha.

The correlation between the test and the retest using the Spearman-Brown coefficient was significant and moderate (.60). When evaluating by condition, we found significant and moderate correlations in the Ori-G (.62) and Self-G (.63) conditions. In the No-G condition, the test-retest correlation was not significant. When evaluating the consistency between test and retest responses with the intraclass correlation coefficient (95% Confidence Interval -CI-), a moderate level of consistency was observed throughout the entire test (ICC = .60, CI: .38-.73). The tests with the Original Glossary (ICC = .63, CI: .22-.83) and with the Self-Generated Glossary (ICC = .61, CI: .19-.82) obtained a moderate level of consistency. The No Glossary test obtained a low level of consistency (ICC = .48, CI: .09-.75). Table 1 summarizes the calculated analyses.

**Table 1.** Internal consistency and test-retest reliability for each condition and the total number of participants using different analyses

|  | Cronbach’s Alpha |  | KR-20 |  | Spearman-Brown | ICC |
| --- | --- | --- | --- | --- | --- | --- |
|  | Test | Retest | Test | Retest |  |  |
| Original Glossary (Ori-G) | .03 | .38 | .06 | .40 | .62* | .63 |
| Self-Generated Glossary (Self-G) | .46 | .60 | .48 | .61 | .63* | .61 |
| No Glossary (No-G) | .23 | .48 | .26 | .50 | .48 | .48 |
| Total | .29 | .51 | .30 | .52 | .60* | .60 |

\* $p < 0.05$ for Spearman-Brown

Additionally, to explore the test-retest reliability, Bland-Altman plots were created for the raw data (Fig 1, Panel A) and for the data transformed to logarithms, because these scores did not meet a normal distribution (Fig 1, Panel B). The entire sample was used in the analysis (n = 89). The average differences were .57 (SD = 3.74) for the raw data and −1.79 (SD = .15) for the data transformed to logarithms. Only two of the 89 points were outside the upper limit of the confidence interval (95%), while the rest were within its limits. High variability in score differences was observed, but the plot with the logarithmic transformation showed that the differences in scores tended to decrease when the average test scores increased.

**Fig 1.**
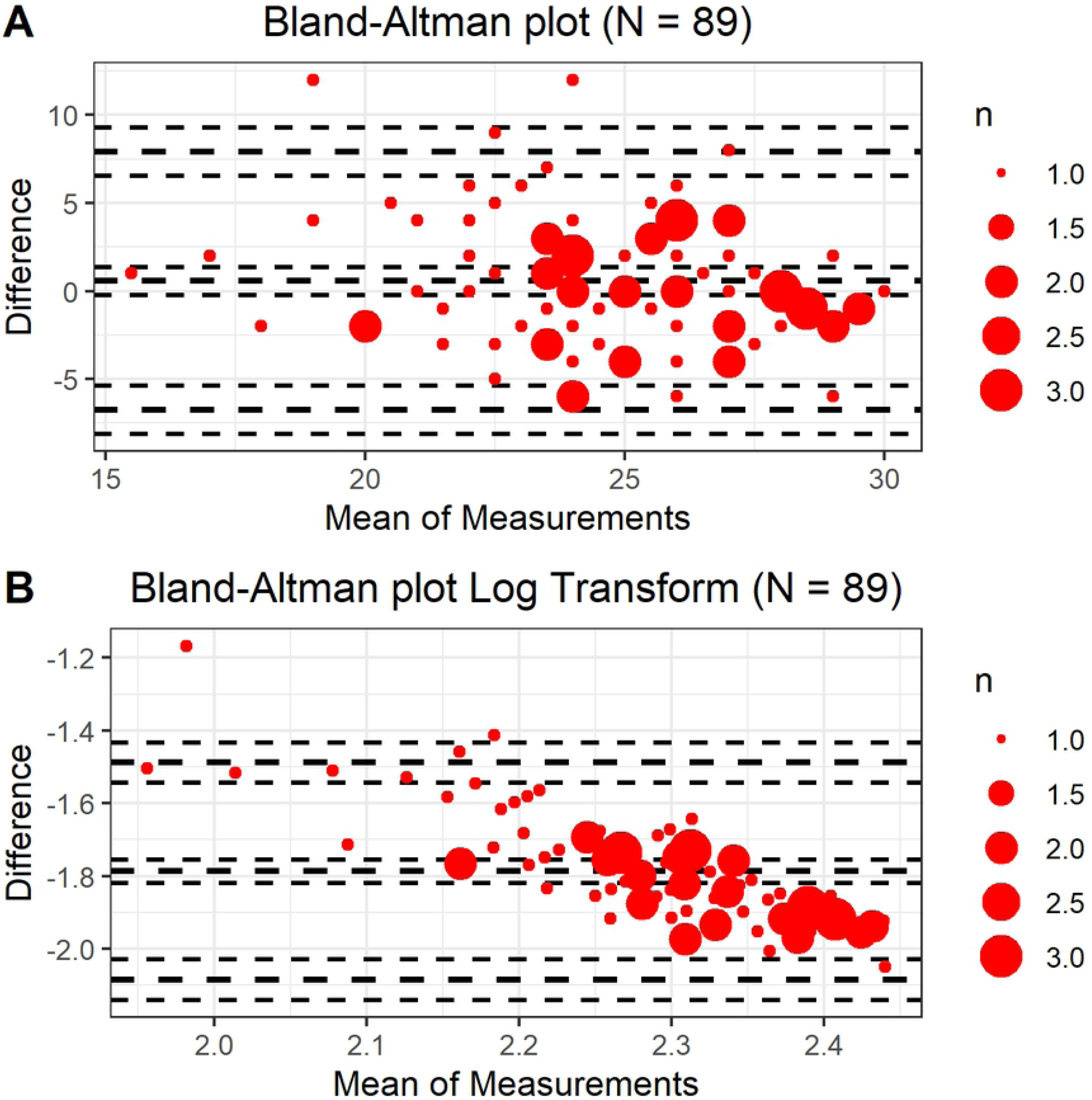
Bland-Altman plot of the differences between test and retest scores. Panel A: Raw scores Panel B: Scores transformed to base 10 logarithms.

### Analysis of average correct answers and average test time

A 3-by-2 mixed ANOVA was implemented with the average number of correct answers. The intra-subject factor consisted of the two times that the test was administered (test and retest), and the inter-subject factor consisted of the three conditions (Original Glossary, Self-Generated Glossary, and No Glossary). There were no significant differences between the three experimental conditions, *F* (2, 86) = 2.28, *p* = .11. In addition, there were no differences between the times the test was administered (Test: M = 25.0, SD = 3.1, Retest: M = 24.4, SD = 3.8), *F* (1, 86) = 2.07, *p* = .15; or the interactions between the instances when the test was administered and the experimental condition, *F* (2, 86) = .13, *p* = .88.

The same 3-by-2 mixed ANOVA was implemented with the average time per stimulus. Significant differences between the conditions were found, 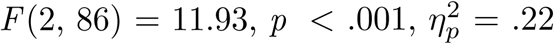. The Self-G (M = 214 *s*) and No-G (M = 217 *s*) conditions did not differ between one another (*p* = .85). However, the Ori-G condition (M = 282 *s*) took significantly longer than the other two conditions (*ps <* .001). Differences were also found between the instances when the test was administered. Regardless of the experimental condition, during the first application of the test, participants took longer (M = 269 s) than in the retest 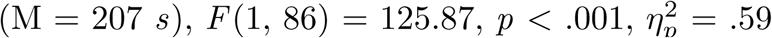. Finally, there was a significant interaction between the conditions and the instances, 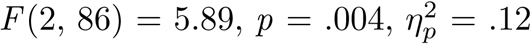. As shown in Fig 2 (Panel B), the time difference between the test and the retest in the Ori-G condition was much greater, 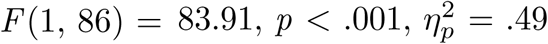, than in the Self-G, 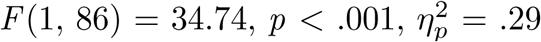; and the No-G conditions, 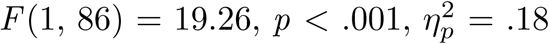.

**Fig 2.**
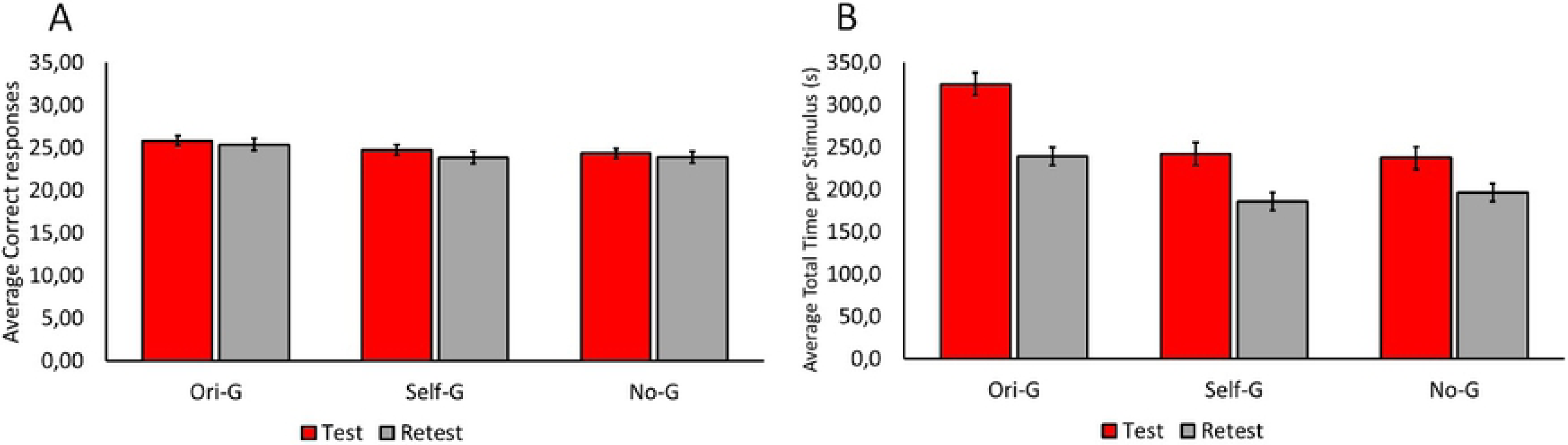
Averages and Standard errors for the average number of correct answers (Panel A) and time taken to complete the test by condition and study phase (Panel B).

## Discussion

The purpose of this research was to determine how the glossary of terms affected the subject’s performance, the temporal stability, and the internal consistency of the Reading the Mind in the Eyes Test. To corroborate this question, we used an experimental manipulation where we generated three different glossary conditions (Original, Self-Generated, and No Glossary). In our study, internal consistency was low, both in the test and in the retest, for all three conditions. These results were similar to those obtained in other studies [17, 20, 24, 28]. In our literature review, we also found broad variability in internal consistency. Comparatively, our study presented the lowest Cronbach’s *α* value of the studies reviewed. Also, even though internal consistency increased in the retest phase, it remained low, considering that a Cronbach’s *α* equal to or greater than .70 is considered adequate [36].

When exploring the internal consistency in each condition, we found that the Ori-G condition had less internal consistency than the other two conditions in which participants created their own glossary (Self-G) and even in the condition in which they did not have a glossary at all (No-G). This trend was observed both the first and second time the test was administered. As shown in Table 1, the Self-G condition was the one that showed the greatest internal consistency.

Like other authors, we found that the test is reliable in terms of temporal stability [10, 15–17]. In our study, the interval between the test and retest was three minutes. Temporal stability did not change between the test administered under the Ori-G and Self-G conditions. However, the temporal stability of the No-G condition was low.

In short, even though both the Self-G and Ori-G conditions had better levels of temporal stability than the No-G condition, the Self-G condition had better levels of internal consistency in both the test and the retest than the Ori-G condition. Therefore, the reliability of the Self-G condition was better than that of the other two conditions in this study.

Preti, Vellante, and Petretto [29] proposed that low reliability reports are due to calculations that violate some assumptions, such as continuity. Hence, they would show inadequate results for a scale with dichotomous responses, such as the RMET. This statement has been refuted by Chalmers [37], who pointed out errors regarding the calculation of reliability and the use of the Ordinal Alpha suggested by Preti, Vellante, and Petretto [29]. Chalmers [37] stated that if the response stimuli are not ordinal, as on a Likert scale, then the Ordinal Alpha is likely inappropriate and should not be used. While tests with dichotomous response items usually have low reliability compared to Likert-type response tests [24], this would not explain the high variability observed with RMET in terms of internal consistency found in various studies, ours among them.

As for the performance of the participants in all three conditions, our average scores were similar to those usually found in the literature. From a sample of twelve studies, our total scores were located between the minimum (M=22.8) and maximum (M=28.4) ranges, and close to the average (M=26.5, SD=1.8) [2, 8, 14, 19, 38–45]. In addition, our study found no significant differences in test and retest scores between the three experimental conditions. Considering that the type of glossary, as well as its presence or absence, did not affect the total number of correct answers, it is apparent that the use of the original glossary or the self-generated glossary did not improve the performance of the participants.

However, participants took less time in the retest than in the test, which can be considered a measure of how well they learned to respond. When exploring the differences by experimental condition, we found that those who built their own glossary and those who did not have a glossary took significantly less time to respond than those in the original glossary condition, both in the test and retest. Since the number of correct answers was similar among the three conditions, our results indicate that it would be better if subjects built their own glossary or didn’t use a glossary at all, considering that participants who used the original glossary took longer to complete the test, yet did no better at responding than the other two conditions.

Our interpretation of the findings is that the condition of creating a Self-Generated Glossary leads to a higher level of processing in the participants [32, 33]. This higher level does not improve participants’ performance, but it improves their velocity and the test’s consistency and stability. This processing gives coherence and stability to the meanings and allows them to be recalled faster. The subjects who were exposed to the original glossary were also able to carry out a level of processing since they reviewed and read the words and meanings in the glossary. However, their level of processing was not very deep because the meanings were learned passively and were not self-generated.

Taking into account the overall results, we suggest the use of the self-generated glossary as an alternative to the original glossary or the absence of a glossary because it achieved greater internal consistency, greater temporal stability, and resulted in a similar number of correct responses compared to the other two conditions. In addition, participants took less time to answer the RMET than those using the original glossary. The absence of glossary did not affect the number of correct answers, although the participants in this condition took the same amount of time to finish the test as those using a self-generated glossary. The problem with this alternative is that while the responses achieved a better level of consistency than the original glossary condition, this procedure had less temporal stability.

One question we have is why our manipulation did not affect the number of correct answers. It is assumed that the RMET measures the ability of participants to match a word with a certain gaze; regulated by meaning-processing mechanisms. However, we propose that the RMET measures a group of abilities that operate independently from each other; related to matching certain gazes with emotions and mental states that the person has learned in the course of his or her social development. Thus, our experimental manipulation, which focused on semantic processing, would affect the temporal stability and consistency of responses, but would not improve the ability to associate a word with a specific gaze.

We recommend interpreting our results of comparisons among experimental conditions with caution, as the reliability obtained was low. While in some cases the reliability of the test was close to satisfactory, the high variability found in different studies remains to be explained. In this regard, we propose using our method of asking participants to build their own glossary. We also recommend exploring the temporal stability of the test with our method using longer time intervals.

We consider that the results obtained in our study are paradoxical. In a strict sense, the test was not consistent even though it was stable. Since the test itself is unreliable, it doesn’t make much sense to evaluate its validity. The scores of our participants, which were quite close to those of adults with normal development, showed no difference among the three experimental conditions, nor between the two instances in which the RMET was applied. We still cannot specify what the test measures. However, we know —based on the levels of stability and consistency— that RMET is sensitive to the experimental manipulations that we carried out.

Finally, we believe it is important to consider that the lack of significant differences in average scores between glossary methods may be because the instrument is not good at detecting differences between individuals with average Theory of Mind ability and a high level of cognitive functioning (Black, 2019).

## Conclusion

The Reading the Mind in the Eyes Test has shown a great ability to differentiate between people with typical development and people with neurocognitive problems. However, some authors have questioned the validity of the test, pointing out that it does not necessarily measure a high ability to detect mental states, mainly because its internal consistency has had broad variability. Our findings indicate that the test has low levels of internal consistency and that the glossary is a potential source of variability. In our case, the higher level of processing we generated experimentally under the Self-Generated Glossary condition did not improve the scores, but made the responses faster, more consistent, and stable.

## Acknowledgments

The authors thank Karen Norambuena for her collaboration during the experiments. This research has been funded by the Comisión Nacional de Investigación Científica y Tecnológica (CONICYT) of the Science, Technology, Knowledge and Innovation Ministry (ANID) under Research Grants REDES-170155 and PCI-PAI80160101 by the Fondo Nacional de Desarrollo Científico y Tecnológico de Chile (FONDECYT) under Research Grant # 1161533, and by the Programa de Investigacioón Asociativa (PIA) en Ciencias Cognitivas de la Universidad de Talca.

